# Revealing Complete Complex *KIR* Haplotypes Phased by Long-read Sequencing Technology

**DOI:** 10.1101/135426

**Authors:** D. Roe, C. Vierra-Green, C.-W. Pyo, K. Eng, R. Kuang, S. Spellman, S. Ranade, D.E. Geraghty, M. Maiers

## Abstract

The killer cell immunoglobulin-like receptor (*KIR*) region of human chromosome 19 contains up to sixteen genes for natural killer (NK) cell receptors that recognize human leukocyte antigen (*HLA*)/peptide complexes and other ligands. The *KIR* proteins fulfill functional roles in infections, pregnancy, autoimmune diseases, and transplantation. However, their characterization remains a constant challenge. Not only are the genes highly homologous due to their recent evolution by tandem duplications, but the region is structurally dynamic due to frequent transposon-mediated recombination. A sequencing approach that precisely captures the complexity of *KIR* haplotypes for functional annotation is desirable.

We present a unique approach to haplotype the *KIR* loci using Single Molecule, Real-Time (SMRT®) Sequencing. Using this method, we have – for the first time – comprehensively sequenced and phased sixteen *KIR* haplotypes from eight individuals without imputation. The information revealed four novel haplotype structures, a novel gene-fusion allele, novel and confirmed insertion/deletion events, a homozygous individual, and overall diversity for the structural haplotypes and their alleles.

These *KIR* haplotypes augment our existing knowledge by providing high-quality references, evolutionary informers, and source material for imputation. The haplotype sequences and gene annotations provide alternative loci for the *KIR* region in the human genome reference GrCh38.p8.

**Author contributions:** **DR, CVG, SS, SR, DEG, MM designed the project. CWP, KE, RH performed the preparation and sequencing experiments. DR, CVG, RK, SR, DWG wrote the majority of the manuscript.**

## Introduction

On human chromosome 19, a 150-250,000 base pair region contains between four and fourteen protein-encoding genes and two pseudo genes of the killer cell immunoglobulin-like receptor (*KIR*) family. These genes encode proteins containing two or three extracellular immunoglobulin-like domains that recognize human leukocyte antigen (*HLA*))/peptide complexes and other ligands^1^. This recognition helps initiate inhibitory or activating cytotoxic signaling in natural killer (NK) cells (and some T cells), utilizing an intracellular immunoreceptor tyrosine-based inhibitory or activation motif (ITIM or ITAM), respectively^2, 3^. NKs and their *KIR* receptors are essential to human health and their genes impact infections (including HIV/AIDs), pregnancy, autoimmune diseases, transplantation, and immunotherapy^3–6^. NK cells induce cytotoxicity against infected or abnormal cells and they release cyto- and chemo-kines as part of a larger immune reaction. This reaction is mediated by competing lack of self antigen recognition and recognition of non-self antigen. Recognition of self inhibits the reaction, and recognition of non-self activates it.

*KIR* genes have duplicated via tandem duplication, deleted, and evolved in primates over the last 30-40 million years^7, 8^. They are 10-15,000 bp long and separated by ∼1,000 bp. Homology is a characteristic of the region, and a pair of alleles of any two genes are approximately 85-98% identical. Another characteristic is structural diversity^9^. The frequency of recombination is high in this system, and dozens of gene-content haplotypes are seen in Europeans alone^10–13^. It is a transposon-rich region which provides the primary mechanism for recurrent meiotic recombination events^13^. The hallmark is a centrally-located 10-15,000 bp region containing a recombination hotspot. This region is an intermediate region between the proximal (conventionally called 'centromeric') region of the haplotype and the distal (or 'telomeric') region. Thus, each full-length haplotype is subdivided into two gene-filled regions separated by an intergenic intermediate region and bound by four ‘framework’ genes: *KIR3DL3* to *KIR3DP1* defining the centromere and *KIR2DL4* to *KIR3DL2* defining the telomere^14^. These centromeric and telomeric regions have frequently recombined around the intermediate hotspot, although some structures are paired more often than others. Recombination can occur between inter- or intra- genic regions, occasionally forming novel functional proteins via fusions from dual genes^13, 15^.

Haplotypes and their centromeric/telomeric regions fall into two categories based on their activating and inhibitory gene content^14^. Class A haplotypes are generally characterized by the presence of inhibitory genes: *KIR2DL4* and *KIR2DS4* are the only gene encoding activating proteins, and most *KIR2DS4* alleles have an exonic deletion which renders them non-functional^16^. Class A haplotypes are structurally invariant (consisting of one gene-content haplotype and its deleted forms) and allelically polymorphic. Class B haplotypes, conversely, are characterized by relatively little allelic polymorphism but greater structural diversity, as they display various combinations of all *KIR* genes. These two classes are in balancing selection, with class A haplotypes generally providing defense against infection and pathogens, and class B haplotypes generally providing reproductive advantages as well as other unknown functions^17^.

Conventionally, *KIR* haplotypes are named as pairs of class A or B regions separated by the intermediate *KIR3DP1-KIR2DL4* intergenic region^14^. The regions are named 'cen' or 'tel', short for 'centromeric' (i.e., proximal) or 'telomeric' (i.e., distal). Unique regions are accessioned with two digits. For example, 'cA01∼tA01' is haplotype consisting of centromeric class A region 01 combined with a telomeric class A region 01. Similarly, 'cB02∼tA01' consists of the centromeric class B region 02 coupled with the telomeric class A region 01. *KIR* haplotype nomenclature is not yet standardized, and this nomenclature follows conventions established by Pyo, et al. in 2010^10^ and adopted by Vierra-Green et al. in 2012^18^.

These characteristics of homology, repetitiveness, and structural diversity have made the region difficult to haplotype. The most comprehensive way to characterize *KIR* haplotypes is to assemble continuous long reads spanning the repetitive sequences comprised within each of the complex haplotypes of a diploid individual. Such completely *de novo* assembled sequences not only provide the ability to discover and annotate *KIR* gene alleles at the highest resolution, but also provide value as references, evolutionary informers, and source material for imputation.

We have demonstrated our ability to overcome these challenges in the sequencing of the entire *KIR* region by comprehensively sequencing haplotypes for eight diploid individuals using a combination of fosmid-based library preparation along and PacBio long-read sequencing. The PacBio® RS II system from Pacific Biosciences was used for sequencing fosmid libraries. These libraries were constructed and screened to obtain complete coverage of both haplotypes for each person. New SMRTbell library preparation approaches were developed for full-length fosmid sequencing to ensure long read lengths which provided unambiguous full haplotype coverage.

## Results

### Haplotype Structures

Sixteen haplotypes from eight individuals were completely and unambiguous sequenced except for two haplotypes whose *KIR3DL3* genes were not captured in the fosmid and a small gap in one of the haplotypes, located in a repetitive insertion spanning over 100,000 bp. The sixteen haplotypes contain nine distinct structures: eight occurrences of the cA01∼tA01 haplotype, and one occurrence each of eight other structures. Figure 1 depicts the nine structures in the context of the four combinations of A/B regional motifs and centromeric/telomeric regions. The lengths range from 69 kb (cA03∼tB02) to 269 kb (cA01∼tB04). Four haplotype structures have not previously been reported: cA04, cB04∼tB03, cB05∼tA01, and cA01∼tB04. Figure 1a shows cA04, a deleted form of cA01∼tA01 containing only the 4 most centromeric genes. Similarly, cA03∼tB02 in Figure 1b depicts a deleted haplotype containing five genes, although the deletion relative to cA01∼tB01 occurred in the middle of the haplotype, leaving the three most centromeric and two most telomeric genes. In contrast, cA01∼tB04 contains a large (>100 kb) insertion of the entire intermediate and tA01 region. Figure 1c depicts cB01∼tB01 and cB04∼tB03, which contains a relative deletion of centromeric *KIR2DS3* through the intermediate and telomeric region up to telomeric *KIR2DL5*. This deletion results in the pairing of *KIR2DL5B\**00201 and *KIR2DS3\**00103. Figure 1d shows the two haplotypes of the cBXX∼tBXX motif from our cohort along with the reference in the middle. Both haplotypes have centromeric deletions relative to the reference.

**Figure 1:**
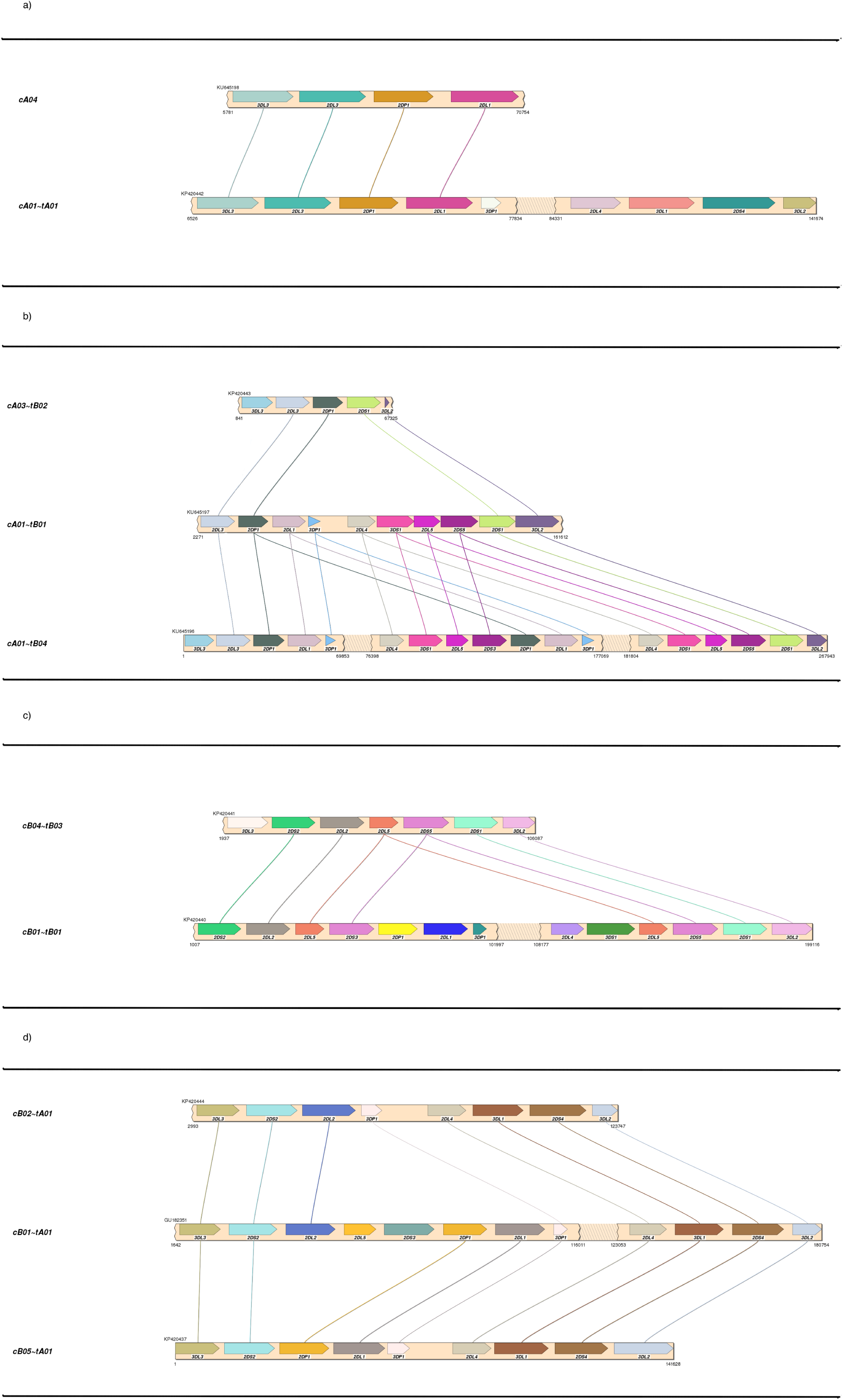
Haplotype structures and regional motifs. Each of the nine structural haplotypes is depicted within the context of pairs of centromeric/telomeric regions and A/B motifs: a) cAXX∼tAXX b) cAXX∼tBXX c) cBXX∼tBXX d) cBXX∼tAXX. cB01∼tA01 was previously published and is included to provide the cBXX∼tAXX reference. Haplotypes cA01∼tB01 and cB01∼tB01 contain *KIR3DL3* but were not captured in the fosmid.

Figure 2 depicts the haplotypes by gene orientation in various contexts. Figure 2a shows that five of the nine haplotype structures contain the four expected framework genes. One haplotype contains six framework genes due to an extra copy of the *KIR3DP1∼KIR2DL4* intermediate region. Three lack *KIR3DP1∼KIR2DL4* due to deletions, one of which also lacks most of its *KIR3DL2* due to the same deletion. Figure 2a also depicts the haplotypes as centromere and telomere regions. Three haplotypes lack the typical full-gene cen∼tel configuration: two have deletions in the telomeric region (cA04 and cA03∼tB02) and one has a duplication in a class B telomeric region (cA01∼tB04). As depicted in Figure 2b, five haplotypes are relative deleted forms and one inserted. Figure 2c breaks down the haplotypes by class A and B regions. Two truncated haplotypes contain no class B (cA04 and cA03∼tB02), while one truncated haplotype does not include class A (cB04∼tB03). One haplotype contains two telomeric regions, both of which are B (cA01∼tB04). The rest break down as two cBxx∼tAxx, one cA01∼tB01, one cA01∼tA01, and one cB01∼tB01. Figure 2d shows the haplotype pair assignments per individual. One individual was found to be base-pair homozygous for cA01∼tA01.

**Figure 2:**
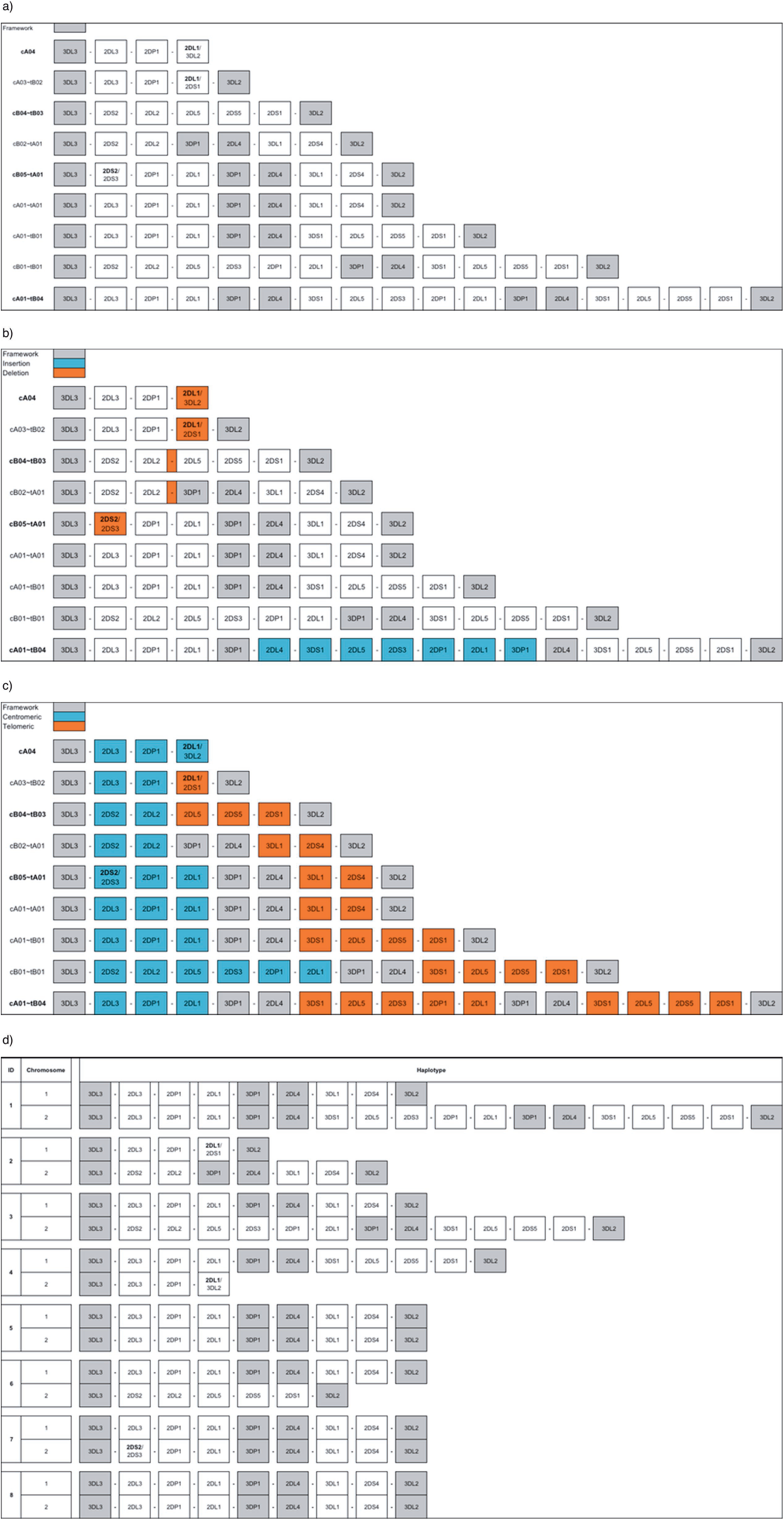
Haplotype structures by gene content. The ordered gene content of the nine distinct haplotype structures are displayed in four categories. a) Framework genes, centromeric and telomeric regions b) Insertion/deletion elements c) Class A and B regions d) Per-individual haplotype pairs. The names of new haplotypes are in bold. Fused genes are depicted with both gene names separated by ‘/’ and the dominant gene name bolded. All haplotypes are completely phased except for one gap in cA01∼tB04, which contains a repetitive insertion spanning over 100,000 bp.

### Novel Fusion Genes and Gene Alleles

Three of the smaller haplotypes contain fusion genes. In Figure 2c, fusion genes are colored yellow, and the gene that gives each fusion its name is in bold. cA04 is a previously unreported haplotype containing a previously unknown fusion between *KIR2DL1* and *KIR3DL2* caused by a deletion of the intermediate and telomeric regions. The new gene consists of *KIR2DL1* 5’ end to intron five and *KIR3DL2* intron six to 3’ end; *KIR2DL1* contains a pseudoexon distal to exon two, and therefore its intron five is structurally equivalent to *KIR3DL2* intron six. The region of recombination is a 129 base pair sequence identically found in both parental alleles. The *KIR2DL1* portion of the sequence is similar to *KIR2DL1\**0020101 except for several intronic SNPs and an exonic SNP at the 13^th^ base that changes phenylalanine to valine. The *KIR3DL2* portion of the sequence is identical to *KIR3DL2*\*0010101. The fusion genes in cA03∼tB02 and cB05∼tA01 have been previously reported^13, 19^: Traherne et al. reported the fusion gene in the same structural haplotype (cA03∼tB02), but the *KIR2DS2/KIR2DS3* fusion gene reported by Rajalingam et al. appears to have been found on a different structural haplotype than in our cohort (cB05∼tA01).

Table 1 reports some high-level characteristics of the gene alleles at several levels of annotation, including alleles of *KIR2DP1* and *KIR3DP1* even though they are pseudogenes. The total number of alleles per gene range from three for *KIR2DL2* and *KIR2DS3* to fourteen for *KIR2DP1* and *KIR3DL2*. Except for the two bordering framework genes of *KIR3DL3* (23% fully sequenced) and *KIR3DL2* (64% fully sequenced), all genes have been characterized comprehensively at the full-gene allele level. Half of the genes contain at least one novel protein-coding allele; the frequency of these novel protein-coding alleles peak at 31% in *KIR2DL1*. *KIR3DL2* is the only gene containing a novel synonymous CDS mutation within a known protein sequence. For nine genes, all full-length alleles were novel. In contrast, all *KIR3DS1* full-length alleles have previously been reported. Alleles of inhibitory proteins, which generally predominate in class A haplotypes, comprise about 60% of the novel alleles at full-gene resolution.

**Table 1:** Novel alleles by gene and resolution. Each of the 16 genes is broken down by frequency of partial alleles and three levels of annotation: protein, CDS, and full-gene. *KIR2DP1* and *KIR3DP1* are pseudogenes.

| **Gene** | **# Alleles** | **% Partial**<br>**sequence** | **% Novel**<br>**protein** | **% Novel CDS** | **% Novel full**<br>**gene** |
| --- | --- | --- | --- | --- | --- |
| *KIR2DL2* | 3 | 0% | 0% | 0% | 100% |
| *KIR2DL3* | 11 | 0% | 0% | 0% | 100% |
| *KIR2DL4* | 13 | 0% | 15% | 15% | 100% |
| *KIR2DL5* | 6 | 0% | 0% | 0% | 100% |
| *KIR2DP1* | 14 | 0% | 50% | 50% | 86% |
| *KIR2DS1* | 5 | 0% | 20% | 20% | 100% |
| *KIR2DS2* | 4 | 0% | 0% | 0% | 100% |
| *KIR2DS3* | 3 | 0% | 0% | 0% | 100% |
| *KIR2DS4* | 9 | 0% | 11% | 11% | 89% |
| *KIR2DS5* | 3 | 0% | 0% | 0% | 100% |
| *KIR3DL1* | 9 | 0% | 22% | 22% | 89% |
| *KIR3DL2* | 14 | 64% | 7% | 14% | 36% |
| *KIR3DL3* | 13 | 23% | 8% | 8% | 77% |
| *KIR3DP1* | 13 | 0% | 69% | 69% | 77% |
| *KIR3DS1* | 4 | 0% | 0% | 0% | 0% |

## Discussion

Our collective knowledge of the *KIR* locus, especially in a translational context, was to date limited by sequencing technologies, with the primary challenge being homology and variation at both large and small scales in the *KIR* locus. In particular, phasing centromeric and telomeric regions across their intermediate region has been impossible due to repetitive homologous sequences spanning well over 10,000 bases. For the first time, we demonstrate the ability to overcome these challenges and provide a complete genetic characterization of full *KIR* haplotypes in a variety of diploid individuals.

Here, we show direct sequencing of linearized full-length fosmids that simplifies the assembly of *KIR* region. We have combined long-read SMRT Sequencing with a fosmid cloning approach. The enrichment of a tiling of targeted fosmids allows cloning 100s of kb of extended lengths of genomic DNA loci of interest. The capability to capture long segments of DNA inserts (∼ 40Kb) and then the sequence them as continuous long-reads provided unique tools necessary for unambiguously assembling two discrete *KIR* haplotypes from each of the diploid samples.

The current maximum read length generated by SMRT Sequencing on PacBio RS II exceeds the length of a full fosmid insert, and it is therefore possible to span an entire fosmid insert with a single continuous read. We have linearized and sequenced complete fosmid inserts to generate a high-quality consensus by simple alignment. In this shotgun sequencing approach, we have removed the need to shear the fosmids before sequencing and assembly. Bypassing the shearing, a step that usually introduced error during shotgun assembly of highly repetitive sequences helped improve the phasing accuracy of the individual fosmid sequences and the high-quality sequences of complete fosmids easily tiled into full haplotypes.

### Full-length Fosmid Sequencing Improved and Augmented Current Haplotype Annotations

Sequencing the fosmid in its entirety is crucial to provide reference haplotypes for interpretation and imputation using lower-resolution typings. Typically, short oligo or primer sequences get used for characterization of protein regions (presence/absence, exon, regulatory), while copy number gets measured via quantitative PCR. On these bases, gene alignment (i.e., haplotypes) and linkage disequilibrium between elements are inferred and applied to association studies. In contrast, complete characterization provides the ability to unite all known functional elements over long ranges for practical considerations and it forms the basis for discovering novel features of all kinds. Haplotype imputation from lower resolution data requires documented structures for correct interpretations. For an example illustrating haplotype information, see Figure 3. A person genotypically homozygous for the A centromeric region and the B telomeric region can only be ambiguously haplotypically imputed. The differences between these ambiguous haplotype pairs can be extensive. For example, the pair of haplotypes could be assigned as homozygous for cA01∼tB01 or cA04 + cA01∼tB04 as seen in Figure 3a. Figure 3b illustrates the sequencing of the transposable elements that caused the deletion resulting in cA04’s *KIR2DL1*/*KIR3DL2* fusion gene: the L2 LINE element is identical in both alleles, with flanking AluSx and MIR SINEs in both alleles. In the top frame, the *KIR2DL1* allele is in the top track and the *KIR3DL2* allele in the bottom; note the track for *KIR2DL1* contains numerous variants on the right side, and the track for *KIR3DL2* contains numerous variants on the left side. See the duplicated region of cA01∼tB04 in Figure 3c; it is 106 thousand base pairs and the two regions are 98% identical.

**Figure 3:**
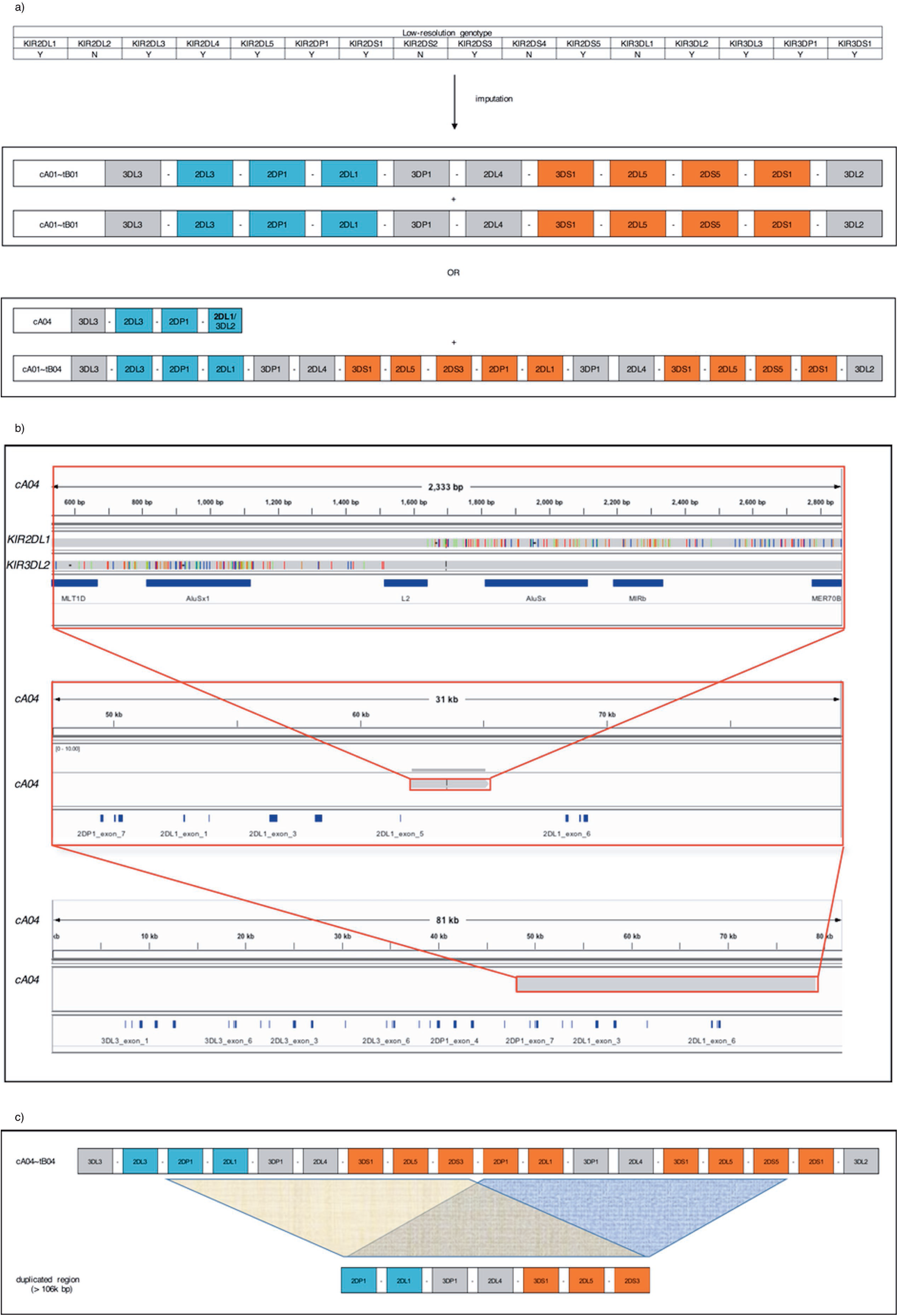
Information gained by fully sequenced and phased haplotypes. a) An individual carrying the given typical low-resolution genotype could be imputed to be homozygous for cA01∼tB01; alternatively, the genotype is imputed as cA04 + cA01∼tB04. b) The transposable elements (dark blue) responsible for the cA04 truncated haplotype, along with their parent alleles from *KIR2DL1* and *KIR3DL2*, are shown in the top frame. The middle frame depicts that same region within the larger. And the bottom frame represents the gene’s location within the haplotype. c) The duplicated region of cA01∼tB04; the two loci are 98% identical over 106 thousand bases.

Our previous multi-resolution genotyping and haplotype inference of these individuals allowed us to assess both the accuracy and quality of these haplotypes. These haplotype sequences are consistent with our previous characterizations^18^ made using multiple lower-resolution time-tested methods such as SSO, SSP, Sanger, TaqMan copy number variation, and physical linkage via haplotype-specific extraction and extended PCR. The only structural difference between our previous haplotype calls is cA04 (*KIR3DL3∼KIR2DL3∼KIR2DP1∼****KIR2DL1****/KIR3DL2*), which was previously reported as *KIR3DL3∼KIR2DL3∼KIR2DS1∼KIR3DL2*; the novel fusion gene was previously unable to be detected and we, therefore, inferred some of the haplotype to be homozygous with its sister chromosome. Twenty-three previously-named alleles have been annotated at the full-gene level, demonstrating the quality of our approach at the base level. Also, our single-molecule, long-read sequencing approach is the only method able to resolve the long (>10kb) repetitive diploid intermediate region between *KIR3DP1* and *KIR2DL4* (Figure 4c); this is crucial to determining haplotype structure due to the recombination hotspot it contains.

We observed increased protein diversity in the inhibitory alleles, consistent with previous reports. We also found a presumably functional novel *KIR2DL1*/*KIR3DL2* fusion gene containing a novel *KIR2DL1-*like protein in the extracellular domain and previously reported *KIR3DL2* intracellular domains.

### The Vast Structural Diversity Provides New Insight into Evolution

The haplotypes range in size from 69,000 base pairs to almost 269,000. The smallest haplotype contains only three non-pseudogenes, and the largest contains an insertion of seven genes over 106,000 base pairs. The annotations in Figure 2 depict the structural characteristics of the region. Overall there is relative structural consistency *within* each of the three regions (centromere, telomere, and intermediate region) and lack of consistency *between* each centromeric or telomeric region, which indicates the relatively frequent recombination *between* the regions as well as the divergent evolution *within* the A and B classes. The region-intermediate hotspot, however, is just one of the locations where transposon-mediated fusions can occur.

Variations beyond binding domains can cause clinical effects. Combinations, alignment, and orientation of genes, regulatory regions, and other non-binding factors such as distance can influence a broad range of phenotypes and provide a framework to interpret functional effects such as expression, licensing, dose effects, etc. The approach we report here is the only method that directly characterizes all these elements of *KIR* genotypes and haplotypes to provide previously unknown reference material, thereby supporting comprehensive genetic annotation and discovery. This study reaffirms the *KIR* region’s extensive structural *and* allelic variation.

## Methods

The Center for International Blood and Marrow Transplant Research (CIBMTR) has a cohort of unrelated individuals with pre-typed information available for *KIR* gene presence/absence and exonic alleles via sequence-specific primer-directed polymerase chain reaction amplification (PCR-SSP), sequence-specific oligonucleotide hybridization (PCR-SSO), and Sanger-sequence based typing (SBT). *KIR* gene content and location in the haplotype was also studied using a combination of haplotype specific extraction, extended PCR, and TaqMan copy number assays. All subjects provided written informed consent for participation in research and the study and consent were approved by the National Marrow Donor Program Institutional Review Board. All the genotypes and structural haplotype predictions were published^18, 20^. This prior information was the basis for the present work, where we studied eight European individuals exhibiting a balance of known/unknown haplotypes, insertion/deletion events, A/B content, and general representation of the diversity of diploid centromeric and telomeric regions.

### Fosmid Library Construction and Screening

Libraries were constructed using vectors and preparative protocols as previously described^12^ at a density of ∼9,000 colonies per well, allowing each library to be distributed over a single 96 well plate. Screening to identify individual wells containing target fosmids for sequencing – content mapping – was carried out using *KIR* haplotyping probes ^21^ as described by the manufacturer (Scisco Genetics Inc., Seattle WA). This approach provided identification of phased clones at two levels, first with probe specificity for *KIR* gene content, and second from the derived sequence data by matching to *KIR* backbone sequences to separate phase. This design allowed for the unambiguous identification of phased fosmid clones spanning all known *KIR* genes and resulting haplotypes.

### Single Fosmid Isolation

Fosmids spanning each *KIR* region were isolated from content-mapped wells using a modification of the recombineering method^22^. The recombineering/targeting probes were designed using ampicillin for selection and locus-specific sequences were derived from regions extending 100 bp within or flanking each *KIR* locus. 17 pairs of 70 bp primers were synthesized (IDT Technologies, Coralville, IA) to amplify targeting probes. Each primer oligo was designed by fusing 50 bp from each half of the targeted sequence with 20 bp from the 5’ and 3’ ends of the ampicillin gene and used in a PCR amplification of the synthesized ampicillin gene (Blue Heron Biotechnology Inc., Bothell, WA). The set of probe sequences listed in Supplementary Table 1 was sufficient to isolate fosmids spanning all of the *KIR* haplotypes described in this report.

Probes selected empirically based on the efficiency of target capture reactions were highly reproducible – library screens routinely provided successful recombineering at a rate of over 98%. Given overlapping fosmid coverage, this rate allowed us to isolate both complete *KIR* haplotypes in a single round of screening from each library.

Pools of targeted *KIR* locus fosmids representing both the *KIR* haplotypes for each of the eight individuals were set up. Full length fosmid insert were generated to prepare a library from 10 μg of each pooled fosmid DNA per sample. Fosmid pools were first treated with Plasmid Safe (Epicentre) at 37°C for 30 minutes to remove any non-specific contaminant bacterial DNA and followed by ampure purification to remove the nuclease enzyme. Fosmids were then linearized to isolate full lengths of the *KIR* locus inserts with Lambda Terminase (Epicentre) using vendor recommended guidelines. The reaction was terminated using ampure purification, instead of heat denaturation, to avoid heat-related DNA damage of the long fosmid template. Additional precaution was taken in handling fosmid DNA during library preparation to prevent DNA damage. The contiguous linear fosmid inserts ready for SMRTbell preparation and were processed using standard Pacific Biosciences 20 Kb library preparation protocol. Approximately 5 μg of SMRTbell library recovered for each of the eight samples size-selected from 15 kb to 50 kb using agarose gel. The size-selected libraries were submitted to an additional DNA Damage Repair at 37°C 60 minutes before SMRT Sequencing on the PacBio RS II platform, using 6 hours movie collection times.

### Sequencing and Data Analysis

Analysis of the full-length fosmid sequences used a modification to the hierarchical genome assembly procedure (HGAP)^23^. The first error correction step of HGAP was carried out using raw reads >15kb aligned against seed reads >25kb. Subsequently, the error corrected reads were filtered for the presence of the flanking fosmid vector and the amp+ selection vector sequences. Within each sequence pool, the number of unique filtered sequences was checked against the expected number of fosmids. If the sequence coverage with full-length fosmid reads was less than the expected number of fosmids, the error corrected reads were re-screened for half-length fosmids, single flanking vectors, and amp+ sequences. The reads were then half-length overlapped to generate full-length fosmids. Once the expected number of full-length error corrected fosmid sequences were generated, they were used to re-map the raw subread data, allowing a high quality ‘quiver’ consensus to be created at >QV50. The amp+ and flanking fosmid vector sequence were trimmed before the fosmids are overlapped using a simple greedy algorithm and a minimum identity of 99.9%^24^.

Genes were annotated relative to IPD-KIR release 2.6.1^25, 26^. Some haplotypes images were generated by SimpleSynteny^27^, using the advanced option.

### Data Access

All sequences have been accessioned in GenBank (KU645195, KU645196, KP420443, KP420444, KP420439, KP420440, KU645197, KU645198, KP420445, KP420446, KU842452, KP420441, KP420438, KP420437, KP420442) and incorporated into GrCh38.p8 (KV575248, KV575249, KV575254, KV575260, KV575246, KV575256, KV575250, KV575257, KV575255, KV575259, KV575247, KV575258, KV575253, KV575252, KV575251).

## Supplementary Data

Supplementary information is available on the *Genes and Immunity* website.

## Acknowledgements

This study was supported by funding from Office of Naval Research grants ONR N00014-12-1- 0142 and ONR N00014-14-1-0028.

We would like to thank Tayebeh Rezaie and Valerie Schneider from the National Center for Biotechnology Information for their assistance incorporating the sequences into the human genome reference, Dan Veltri for help and advice with SimpleSynteny, and Julia Udell and Michael Wright for reviewing the manuscript.

## References

1 Rajalingam R. Human diversity of killer cell immunoglobulin-like receptors and disease. Korean J Hematol 2011; 46: 216.

2 Vilches C, Parham P. KIR: Diverse, Rapidly Evolving Receptors of Innate and Adaptive Immunity. Annu Rev Immunol 2002; 20: 217–251.

3 Parham P, Moffett A. Variable NK cell receptors and their MHC class I ligands in immunity, reproduction and human evolution. Nat Rev Immunol 2013; 13: 133– 144.

4 Foley B, Felices M, Cichocki F, Cooley S, Verneris MR, Miller JS. The biology of NK cells and their receptors affects clinical outcomes after hematopoietic cell transplantation (HCT). Immunol Rev 2014; 258: 45–63.

5 Davis C, Rizzieri D. Immunotherapeutic Applications of NK Cells. Pharmaceuticals 2015; 8: 250–256.

6 Carrington M, Nelson GW, Martin MP, Kissner T, Vlahov D, Goedert JJ et al. HLA and HIV-1: heterozygote advantage and B*35-Cw*04 disadvantage. Science 1999; 283: 1748–1752.

7 Martin AM, Freitas EM, Witt CS, Christiansen FT. The genomic organization and evolution of the natural killer immunoglobulin-like receptor (KIR) gene cluster. Immunogenetics 2000; 51: 268–280.

8 Carrillo-Bustamante P, Keşmir C, de Boer RJ. The evolution of natural killer cell receptors. Immunogenetics 2016; 68: 3–18.

9 Martin AM, Kulski JK, Gaudieri S, Witt CS, Freitas EM, Trowsdale J et al. Comparative genomic analysis, diversity and evolution of two KIR haplotypes A and B. Gene 2004; 335: 121–131.

10 Pyo C-W, Guethlein LA, Vu Q, Wang R, Abi-Rached L, Norman PJ et al. Different Patterns of Evolution in the Centromeric and Telomeric Regions of Group A and B Haplotypes of the Human Killer Cell Ig-Like Receptor Locus. PLoS ONE 2010; 5: e15115.

11 Jiang W, Johnson C, Jayaraman J, Simecek N, Noble J, Moffatt MF et al. Copy number variation leads to considerable diversity for B but not A haplotypes of the human KIR genes encoding NK cell receptors. Genome Res 2012; 22: 1845– 1854.

12 Pyo C-W, Wang R, Vu Q, Cereb N, Yang SY, Duh F-M et al. Recombinant structures expand and contract inter and intragenic diversification at the KIR locus. BMC Genomics 2013; 14: 89.

13 Traherne JA, Martin M, Ward R, Ohashi M, Pellett F, Gladman D et al. Mechanisms of copy number variation and hybrid gene formation in the KIR immune gene complex. Hum Mol Genet 2010; 19: 737–751.

14 Hsu KC, Chida S, Geraghty DE, Dupont B. The killer cell immunoglobulin-like receptor (KIR) genomic region: gene-order, haplotypes and allelic polymorphism. Immunol Rev 2002; 190: 40–52.

15 Ordóñez D, Gómez-Lozano N, Rosales L, Vilches C. Molecular characterisation of KIR2DS2*005, a fusion gene associated with a shortened KIR haplotype. Genes Immun 2011; 12: 544–551.

16 Graef T, Moesta AK, Norman PJ, Abi-Rached L, Vago L, Older Aguilar AM et al. KIR2DS4 is a product of gene conversion with KIR3DL2 that introduced specificity for HLA-A*11 while diminishing avidity for HLA-C. J Exp Med 2009; 206: 2557–2572.

17 Manser AR, Weinhold S, Uhrberg M. Human KIR repertoires: shaped by genetic diversity and evolution. Immunol Rev 2015; 267: 178–196.

18 Vierra-Green C, Roe D, Hou L, Hurley CK, Rajalingam R, Reed E et al. Allele-Level Haplotype Frequencies and Pairwise Linkage Disequilibrium for 14 KIR Loci in 506 European-American Individuals. PLoS ONE 2012; 7: e47491.

19 Rajalingam R, Gardiner CM, Canavez F, Vilches C, Parham P. Identification of seventeen novel KIR variants: fourteen of them from two non-Caucasian donors. Tissue Antigens 2001; 57: 22–31.

20 Vierra-Green C, Roe D, Jayaraman J, Trowsdale J, Traherne J, Kuang R et al. Estimating KIR Haplotype Frequencies on a Cohort of 10,000 Individuals: A Comprehensive Study on Population Variations, Typing Resolutions, and Reference Haplotypes. PLOS ONE 2016; 11: e0163973.

21 Nelson WC, Pyo C-W, Vogan D, Wang R, Pyon Y-S, Hennessey C et al. An integrated genotyping approach for HLA and other complex genetic systems. Hum Immunol 2015. doi:10.1016/j.humimm.2015.05.001.

22 Nedelkova M, Maresca M, Fu J, Rostovskaya M, Chenna R, Thiede C et al. Targeted isolation of cloned genomic regions by recombineering for haplotype phasing and isogenic targeting. Nucleic Acids Res 2011; 39: e137–e137.

23 Chin C-S, Alexander DH, Marks P, Klammer AA, Drake J, Heiner C et al. Nonhybrid, finished microbial genome assemblies from long-read SMRT sequencing data. Nat Methods 2013; 10: 563–569.

24 Kearse M, Moir R, Wilson A, Stones-Havas S, Cheung M, Sturrock S et al. Geneious Basic: An integrated and extendable desktop software platform for the organization and analysis of sequence data. Bioinformatics 2012; 28: 1647– 1649.

25 Robinson J, Halliwell JA, Hayhurst JD, Flicek P, Parham P, Marsh SGE. The IPD and IMGT/HLA database: allele variant databases. Nucleic Acids Res 2014. doi:10.1093/nar/gku1161.

26 Marsh SGE, Parham P, Dupont B, Geraghty DE, Trowsdale J, Middleton D et al. Killer-cell immunoglobulin-like receptor (KIR) nomenclature report, 2002. Immunogenetics 2003; 55: 220–226.

27 Veltri D, Wight MM, Crouch JA. SimpleSynteny: a web-based tool for visualization of microsynteny across multiple species. Nucleic Acids Res 2016; 44: W41–W45.

